# The Phospholipase A2 Activity of Peroxiredoxin 6 Protects Hepatocarcinoma Cells Against Ferroptosis and Mitophagy Impairment

**DOI:** 10.64898/2026.11.06.764823

**Authors:** Ángel Ortiz-Alcántara, Alberto Ortiz-Olivencia, Raúl González-Ojeda, Brian McDonagh, Raquel Requejo-Aguilar, C. Alicia Padilla

## Abstract

Peroxiredoxin 6 (PRDX6) is a moonlighting protein that plays a key role in regulating redox homeostasis and lipid metabolism through its unique dual peroxidase and calcium-independent phospholipase A2 (iPLA2) activities. Here, we generated a stable HepG2 cell line harboring a D140A mutation to selectively ablate iPLA2 activity, and we compared its phenotype with that of wild-type (WT) and PRDX6-knockout (KO) cells. D140A exhibited low ROS leves, similar to WT cells, confirming that oxidative stress is due to peroxidase deficiency. In contrast, D140A cells displayed higher lipid peroxidation than WT cells, though significantly lower than that of KO cells. Furthermore, the D140A mutant mirrored the KO phenotype regarding intracellular ferrous iron accumulation, impaired lipid droplet biogenesis, and the downregulation of the key ferroptosis regulator GPX4. Mechanistically, integrated proteomic, lipidomic, and enzymatic analyses revealed that a deficiency in iPLA2 activity of PRDX6 disrupted the release of key lipid species necessary for cell structure and signaling. Consequently, organelles involved in lipid synthesis and trafficking, such as peroxisomes, endoplasmic reticulum, mitochondria and lipid droplet, were affected. Specifically, we found that mitochondria exhibited a reduction in cristae and a diminished capacity for mitophagy and autophagy, which ultimately makes cells much more susceptible to erastin-induced ferroptotic cell death. Taken together, our findings demonstrate that PRDX6’s iPLA2 activity uniquely coordinates organelle lipid homeostasis and mitophagy to protect cancer cells from ferroptotic cell death. Consequently, the use of iPLA2 inhibitors such as MJ33, which decreased PRDX6 iPLA2 activity to a level comparable to the mutant D140A, could represent a promising therapeutic strategy for inducing ferroptotic cell death in hepatocellular carcinoma.

## INTRODUCTION

Peroxiredoxin 6 (PRDX6) is a moonlighting protein that plays a key role in regulating redox homeostasis and lipid metabolism. Distinct from other PRDXs, it uniquely acts as a peroxidase on phospholipid hydroperoxides ^1^ and possesses two additional activities: Ca^2+^-independent phospholipase A2 (iPLA2) ^2^ and lysophosphatidylcholine acyltransferase (LPCAT) ^3^. This implies that PRDX6 is involved in lipid membrane repair and turnover by eliminating peroxidized lipids. Furthermore, PRDX6 is the only peroxiredoxin capable of translocating to damaged mitochondrial membranes, regulating mitophagy by suppressing ROS ^4^. Consistent with these roles, previous studies have demonstrated that PRDX6 deficiency in hepatocarcinoma SNU475 cells elevates ROS production, induces mitochondrial alterations, enhances lipid peroxidation, and alters lipid droplet formation, ultimately leading to increased susceptibility of these tumor cells to ferroptosis ^5^.

Ferroptosis is a form of regulated cell death driven by uncontrolled, iron-dependent lipid peroxidation, causing terminal damage to the membrane and subsequent cell death ^6^. It has become a key area in redox biology and a promising strategy for treating human diseases ^7–9^. In particular, its effectiveness against therapy-resistant and metastatic cancers makes it a highly attractive therapeutic target ^10–13^. Ferroptosis is primarily governed by three specific key factors: the failure to repair lipid peroxide, the oxidation of polyunsaturated fatty acid (PUFA)-containing phospholipids, and the presence of redox-active iron ^14^. Cellular membranes are rich in phospholipids that are incorporated with PUFAs, making them highly susceptible to peroxidation. The peroxidation of PUFAs-containing phospholipids in membranes compromises membrane integrity, increases permeability, and leads to membrane rupture and cell death ^15^. Elucidating the distribution of iron and the formation of oxidized lipids are pivotal to understanding the mechanisms underlying ferroptosis induction. Recent studies demonstrate that while lipid peroxidation (LPO) within diverse subcellular membranes can trigger ferroptosis, the endoplasmic reticulum (ER) represents a critical site for this process ^16^. Furthermore, intra-lysosomal LPO induces lysosomal membrane permeabilization (LMP), which prompts subsequent iron leakage; this cascade accelerates cell-wide LPO, ultimately driving ferroptotic cell death in tumor cells ^17^. The synthesis and remodeling of these phospholipids is primarily driven by two key players: acyl-CoA synthetase long-chain family member 4 (ACSL4) and lysophosphatidylcholine acyltransferase 3 (LPCAT3) that catalyze the reacylation of lysophospholipids. This process generates PUFA-containing phospholipids predominantly PC and PE, in the ER ^18–21^. Evidence indicates that elevated levels of ACSL4 and LPCAT3 cause cells to accumulate more polyunsaturated fatty acid (PUFA) phospholipids, thereby increasing their susceptibility to ferroptosis driven by lipid peroxidation under oxidative stress ^22,23^. These PUFA-rich phospholipids are essential for maintaining the fluidity and flexibility of the inner and outer mitochondrial membranes ^24,25^, but at the same time, excess unesterified PUFAs are highly lipotoxic. Elevated lipid peroxidation in mitochondria leads to severe structural breakdown of the inner mitochondrial membrane, initially triggering mitophagy. However, sustained structural failure and the loss of healthy lipid buffering systems can compromise mitochondrial turnover. This can also cause altered expression of mitochondrial chain complex subunits, resulting from progressive membrane degradation. In this context, attempts to induce autophagy to clean up damaged organelles are ultimately blocked by the accumulation of toxic lipid species. Failure to process and clear autophagosomes causes lysosomal dysfunction. Rather than promoting survival, the impaired autophagic system becomes a pro-death mechanism that accelerates tumour cell collapse via integrated ferroptotic and autophagic stress pathways ^26^.To prevent this, cells convert PUFAs into neutral lipids, primarily triglycerides (TGs), and store them inside lipid droplets (LDs) ^25,27^, which form close physical contact sites with mitochondria to act as central hubs for lipid trafficking. Through these junctions, stored PUFAs can be precisely channeled to the mitochondria for energy production and membrane integrity ^28–31^. The dynamic turnover of LDs constitutes an essential adaptive response in stressed cellular environments. Consequently, elevated LD production acts as a highly conserved cytoprotective mechanism; numerous studies demonstrate that impaired LD assembly under stress compromises viability and triggers cell death ^32,33^. Furthermore, LDs can also accumulate peroxidized lipids, which conversely, could predispose cells to ferroptotic death ^34,35^.

Regulatory networks, such as the p62-Keap1-NRF2 pathway, integrate cellular stress signals to modulate ferroptosis sensitivity ^36^. It has been reported that resistance to ferroptosis in hepatocellular carcinoma cells is mediated by this p62-Keap1-NRF2 pathway, which upregulates key genes involved in iron and ROS homeostasis, such as quinone oxidoreductase 1 (NQO1), heme oxygenase-1 (HMOX1), and ferritin heavy chain 1 (FTH1) ^37^. Under oxidative stress, p62 sequesters Keap1, allowing nuclear factor erythroid 2-related factor 2 (NRF2) to translocate to the nucleus and activate antioxidant and iron-regulatory genes. NRF2 upregulates enzymes involved in GSH synthesis (e.g., GCLC, GCLM), GPX4, and HMOX1) while simultaneously enhancing iron storage (via ferritin) and export (via ferroportin) ^38^. This dual regulation mitigates both oxidative damage and labile iron accumulation, highlighting NRF2 as a regulator of ferroptosis resistance.

Among the enzymatic defense systems that detoxify harmful phospholipid hydroperoxides (PLOOH) to prevent ferroptosis, selenium-dependent glutathione peroxidase 4 (GPX4) was the first to be described and the most extensively studied ^39–41^. However, it has been discovered that there are other regulatory pathways of ferroptosis that are independent of GPX4 ^42^. These include the NAD(P)/FSP1/ubiquinone or vitamin K axis ^43,44^, the GCH1/dihydrobiopterin/DHFR system ^45,46^ and DHCR ^47^, all of which can compensate for GPX4 loss in specific cell types. In contrast to GPX4, these pathways inhibit the propagation of lipid peroxidation by converting phospholipid peroxyl radicals into intermediaries that undergo further decomposition. However, like GPX4, PRDX6 enzyme is capable of directly reducing PLOOHs to their corresponding alcohols and thus plays a key role in protecting against ferroptosis ^5,48–50^. Still, the mechanism by which PRDX6, or one of its activities, regulates susceptibility to ferroptosis or its impact on PLOOH reduction is not well understood. Recent research highlights two key roles by which PRDX6 interact with GPX4 during ferroptosis. First, PRDX6 functions as a selenium trafficking required for the biosynthesis of essential selenoproteins such as GPX4 ^51–53^. Second, it serves as a key as modulator of GPX4 by regulating its cellular localization and activity, ultimately promoting resistance to ferroptosis ^54^.

In this study, a PRDX6 iPLA2 activity deficient mutant was generated from HepG2 hepatocarcinoma cells. A comparative analysis was subsequently conducted to investigate the role of this enzymatic activity in the lipid remodeling, involving alterations and crosstalk among lysosomes, peroxisomes, endoplasmic reticulum, mitochondria and lipid droplet. Ultimately, these interconnected changes culminate in the blockade of mitophagy and autophagy, thereby sensitizing these cells to ferroptotic cell death.

## RESULTS

### Generation of a monoclonal HepG2 cell line expressing PRDX6 lacking iPLA2 activity (HepG2D140A)

The PLA2 activity of PRDX6 functions to hydrolyze phospholipids at the sn-2 position to produce free fatty acids and lysophospholipids. Unlike most PLA2 enzymes, this activity is calcium-independent and has a strict catalytic triad (S32, H26, and D140). Within PRDX6, H26 binds to the liposomal surface, S32 stabilizes the protein structure and binds the phospholipid substrate, and D140 is involved in catalysis ^55^. In this study, a stable HepG2 cell line with the D140A mutation was established, which consequently impairs the iPLA2 activity of PRDX6.

A PRDX6-deficient (KO) HepG2 cell line, generated using CRISPR/Cas9 ^56^, was used as the starting point. These cells were transfected with the pcDNA-*prdx6D140A* plasmid, which expressed a PRDX6 protein lacking iPLA2 activity, and subjected to a selection process using H_2_O_2_. It had been observed that PRDX6KO cells were more sensitive to H_2_O_2_ than WT cells (Figure 1A), largely due to the lack of peroxidase activity; therefore, transfected cells would be more resistant to H_2_O_2_ treatment, since they possess peroxidase activity. Following this selection, limiting dilution was performed to obtain a stable monoclonal cell line lacking iPLA2 activity. Several positive clones were obtained, and the clone exhibiting the highest levels of PRDX6 expression was selected. This expression was determined by Western blot and immunofluorescence using specific antibodies for PRDX6 and was approximately 70% of that expressed by HepG2WT cells (Figure 1B). Furthermore, the presence of the D140A mutation in the mutant cell line was confirmed by Sanger sequencing (Figure 1C).

**Figure 1.**
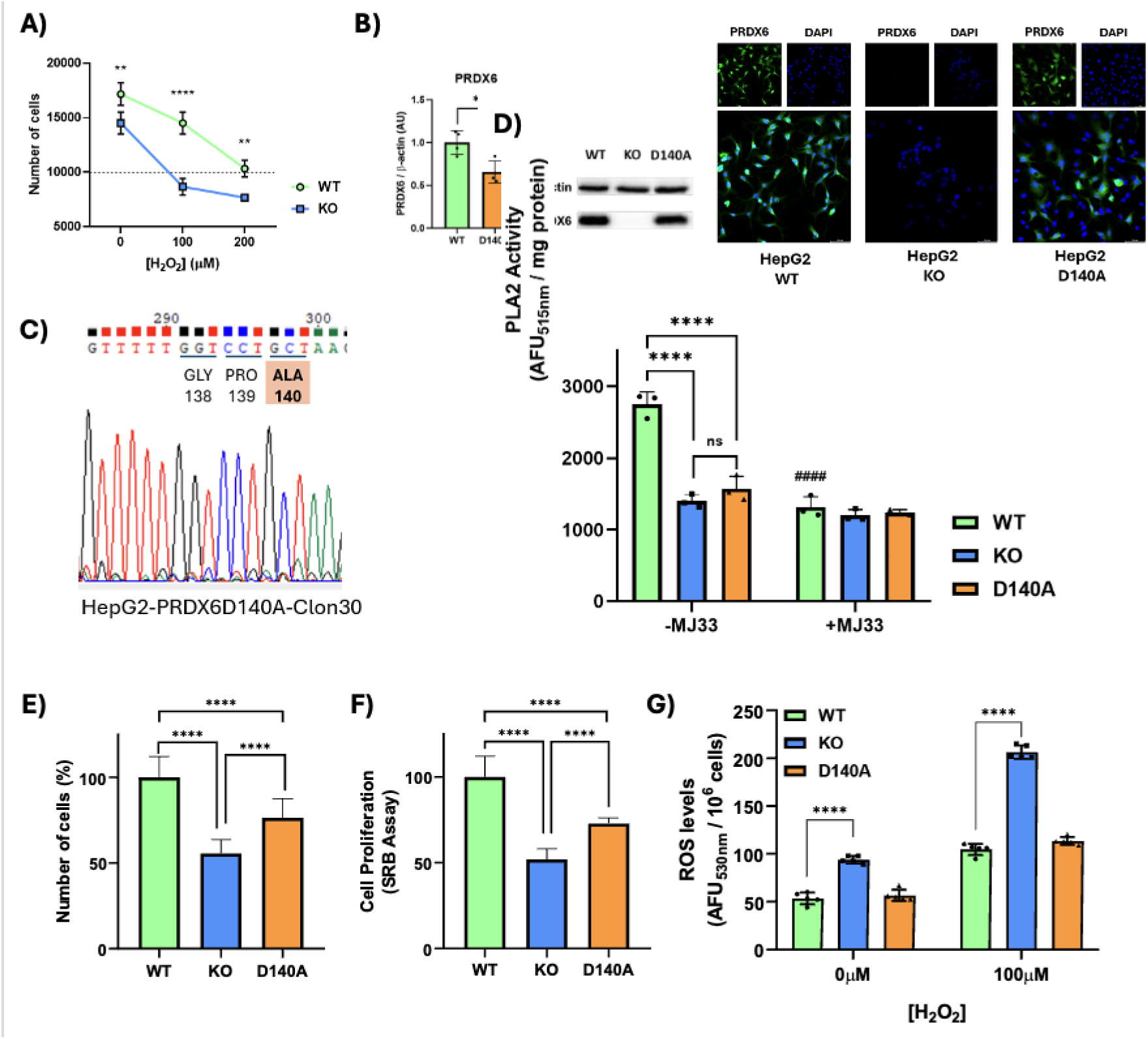
Construction and characterization of an iPLA2-deficient PRDX6 (D140A) HepG2 cell line, in which cell proliferation is impaired without an increase in oxidative stress. (**A**) Growth curves of H₂O₂ dose-response in HepG2WT and HepG2KO cells were used to determine the selection concentration for enriching KO cells transfected with the pcDNA3.1 plasmid containing the mutant PRDX6D140A. (**B**) Mutant D140A expression was validated by Western-blot and immunofluorescence. (**C**) DNA sequencing of the HepG2D140A, showing the presence of the mutation. (D) PLA2 activity exhibited in HepG2WT, KO and D140A cell lines treated or not with MJ33 10 μM. (**E**) Relative cell number (%) in HepG2WT, KO and D140A cell lines. (**F**) Cell proliferation assessed by SRB assay in HepG2WT, KO and D140A cell lines. (**G**) Intracellular ROS levels HepG2WT, KO and D140A cell lines at 60 minutes under basal conditions and following treatment with H₂O₂ 100 μM. Student’s *t*-test (**A, B**), One-way ANOVA (**E, F, G**) and Two-way ANOVA (**D**) analysis were performed. Significance levels: \**P* < 0.05; \*\**P* < 0.01; \*\*\*\**P* < 0.0001.

In addition, iPLA2 activity was determined in extracts from the three cell types under study using the assay described in “Materials and Methods”. iPLA2 activity was significantly lower in HepG2KO and D140A mutant cells compared directly to the WT control group (Figure 1D). Crucially, this suppressed enzymatic activity in KO and D140A cells was statistically comparable to the baseline levels observed when HepG2WT extract cells were treated with MJ33, a specific inhibitor of iPLA2 activity.

### The effect of PRDX6 on cell proliferation is mediated by both its peroxidase and phospholipase A2 activities

It has been reported in HepG2 and SNU475 hepatocarcinoma cell lines, as well as in HCT116 colon cancer cell line, that the complete absence of PRDX6 produces an inhibitory effect on cell proliferation ^56–58^. In this study, proliferation was significantly reduced in both KO and D140A mutant cells compared to HepG2WT cells; however, the former displayed a 50% decrease whereas the latter showed a minor reduction of 25%. This result demonstrated that both activities were necessary for PRDX6 to exert an effect on this process, which is known to be crucial during tumorigenesis. Cell proliferation was determined both by trypan blue cell counting (Figure 1E) and by SRB staining (Figure 1F). This result is highly consistent with the multifaceted and tightly controlled regulation of the cell cycle and cell proliferation. These cellular processes are driven by complex intracellular signaling cascades that are heavily influenced and modulated by both ROS homeostasis and lipid-mediated signaling pathways which are directly linked to peroxidase and iPLA2 activities of PRDX6.

### The increased ROS levels in PRDX6-depleted HepG2 cells are independent of iPLA2 activity

It had previously been observed that ROS levels in HepG2 and SNU475 hepatocarcinoma cells, as well as in HCT116 colon cancer cells, were significantly higher in cells lacking PRDX6. Here, ROS levels were determined in WT, KO, and D140A HepG2 cells, under both basal conditions and oxidative stress induced by H₂O₂ treatment. Conversely, ROS levels in D140A mutant cells remained comparable to those observed in WT cells. Therefore, these results demonstrate that the peroxidase activity of PRDX6, rather than its phospholipase A2 activity, is strictly required for ROS scavenging in this experimental setting (Figure 1G).

### The ablation of PDX6 iPLA2 activity sensitizes HepG2 cells to ferroptosis via iron and peroxidized lipid accumulation with reduction of GPX4 expression

Recent studies have demonstrated the role of PRDX6 in protecting against ferroptotic cell death and its functional relationship with GPX4 ^5,53,59,60^. It remains controversial whether it acts independently or alongside GPX4, and which of its enzymatic activities is driving this process.

Here, we conducted a comparative study among the three HepG2 cell lines (WT, KO, and D140A) to analyze their sensitivity to the ferroptosis-inducing agent, erastin, as well as the rescue effect of ferrostatin-1. In HepG2KO and D140A mutant cells, a deficiency in iPLA2 activity increased vulnerability to ferroptotic cell death compared to HepG2WT cells when exposed to ferroptosis inducer such as erastin, an inhibitor of the xCT cystine/glutamate antiporter (Figure 2A). This increased sensitivity to ferroptosis was reversed by treatment with ferrostatin-1, a selective inhibitor of ferroptosis. These effects were determined via cell counting. The percentage of KO and D140A cells following erastin treatment, relative to their respective controls, was very similar (56% and 58%, respectively), indicating that the phospholipase A2 activity of PRDX6 was what mediated protection against ferroptosis.

**Figure 2.**
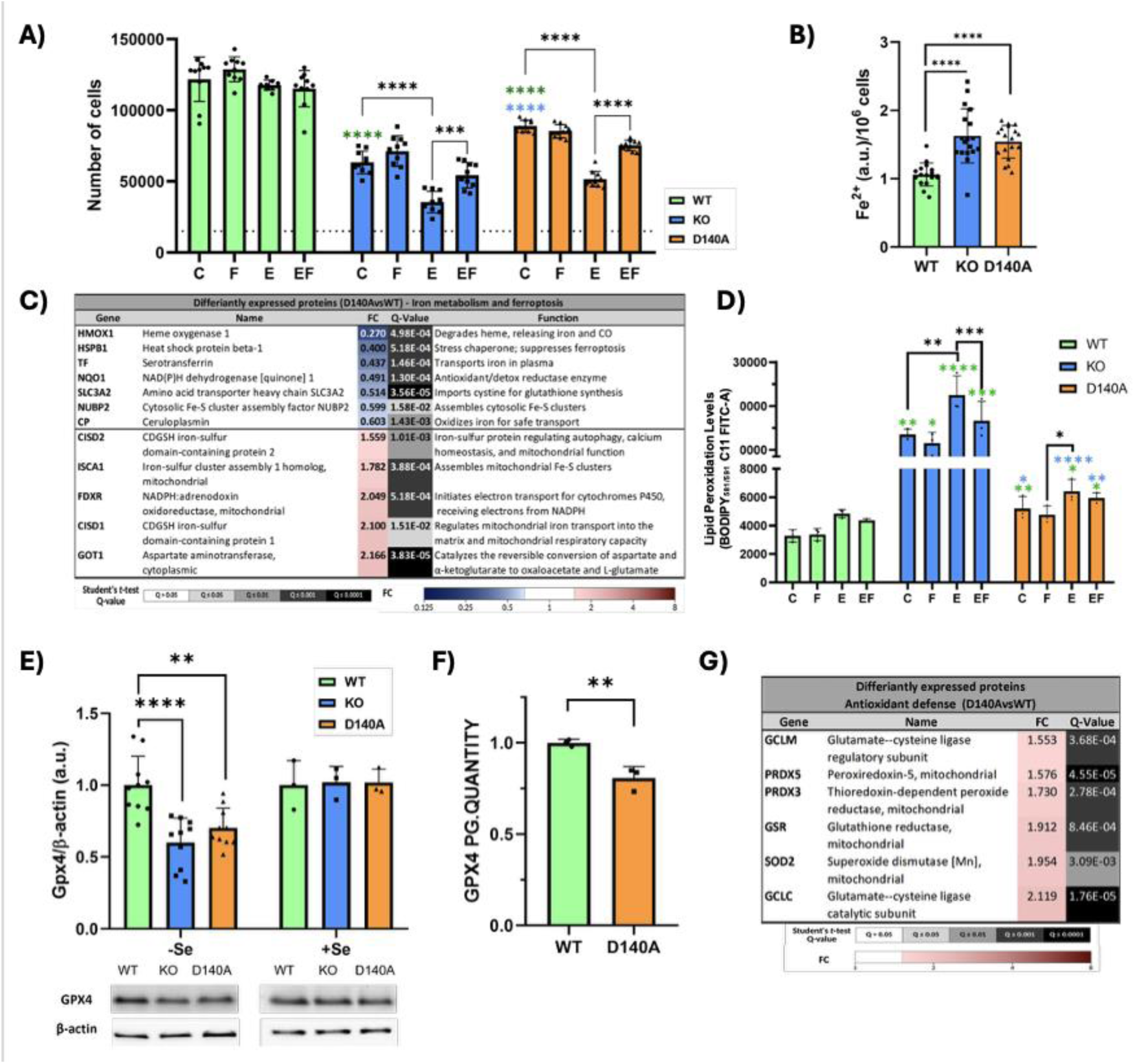
Loss of PRDX6 iPLA2 activity sensitizes HepG2 cells to erastin-induced ferroptosis through dysregulation of iron homeostasis, an increase in lipid peroxidation and a reduction in GPX4 levels and triggering a compensatory antioxidant response. (A) Cell number in HepG2WT, KO and D140A cell lines under control (C), ferrostatin-1 10 μM (F), erastin 4 μM (E), and combined (EF) treatments. (B) Intracellular Fe²⁺ levels in HepG2WT, KO and D140A cell lines. (C) Differentially expressed proteins related to iron metabolism and ferroptosis (D140A vs. WT) identified by proteomic analysis. (D) Lipid peroxidation levels quantified by BODIPY C11 581/591 molecular probe in HepG2WT, KO and D140A cell lines across the same treatment conditions. (E) GPX4 protein expression assessed by Western-blot in HepG2WT, KO and D140A cells cultured with (+Se) or without (-Se) sodium selenite 100nM. (F) GPX4 protein levels quantified by proteomic analysis in HepG2WT and D140A cells. (G) Differentially expressed proteins related to antioxidant defense (D140A vs. WT) identified by proteomic analysis. Student’s *t*-test (C, E, F, G), One-way ANOVA (B) and Two-way ANOVA (A, D) statistical analyses were performed. Significance levels: \**P* < 0.05; \*\**P* < 0.01; \*\*\**P* < 0.001; \*\*\*\**P* < 0.0001.

During ferroptosis, the intracellular labile iron expanded significantly. This excess Fe²⁺ drives the Fenton reaction, interacting with hydrogen peroxide to generate highly reactive hydroxyl radicals (⋅ OH). These radicals strip electrons from PUFAs in cell membranes, initiating a deadly chain reaction of lipid peroxidation.

We have found that intracellular iron levels increased significantly in HepG2KO cells compared to WT, while the D140A mutant behaved similarly to the KO, suggesting this specific mutation leads to the same iron dysregulation (Figure 2B). The results obtained through proteomic analysis corroborated this alteration in cells lacking PRDX6 iPLA2 activity. The simultaneous decrease of HMOX1, HSPB1, TF, NQO1, SLC3A2, NUBP2, and CP (Figure 2C) points toward an alteration in cellular iron transport, and a potential vulnerability to ferroptosis, since low levels of transferrin (TF), ceruloplasmin (CP), nucleotide binding protein 2 (NUBP2) and hemo oxigenase 1 (HMOX1) alter cellular iron trafficking and the intracellular labile iron pool. On the other hand, the reduction of the SLC3A2 limits cystine uptake. This directly impairs glutathione synthesis, thereby compromising the cellular capacity to mitigate lipid peroxidation mediated by enzymes such as GPX4 or PRDX6. Peroxidized lipids were also determined, with a significantly greater accumulation observed in HepG2D140A than in WT (Figure 2D) but much less than in the HepG2KO cell line.

In addition, GPX4 was significantly reduced in both KO and D140A mutant cells compared to WT. This was determined by Western blot (Figure 2E) and proteomic analyses (Figure 2F), with both results being consistent. Additionally, GPX4 expression increased when HepG2 cells were cultured with selenium. This effect was independent of PRDX6, meaning that selenium can upregulate GPX4 even without the presence of PRDX6 (Figure 2E).

The lack of PRDX6 iPLA2 activity drastically compromised membrane lipid homeostasis, sensitizing the cells to iron and lipid peroxidation dependent ferroptosis. To counteract this, the cells deployed an emergency response, characterized by a marked upregulation of key antioxidant enzymes. Specifically, GCLC, GCLM, and GSR were induced to enhance glutathione synthesis and recycling, while SOD2, PRDX3, and PRDX5 reinforced mitochondrial radical scavenging (Figure 2G). Additionally, expression of five anti-ferroptotic proteins (GOT1, CISD1/2, FDXR, and ISCA1) was significantly increased (Figure 2C) ^61–65^.

Consequently, HepG2 cells expressing the PRDX6D140A mutant remain viable and proliferate, although to a lesser extent than WT cells. Furthermore, they exhibit increased susceptibility to ferroptosis when challenged with inducers such as erastin.

### The absence of PRDX6 iPLA2 activity impairs mitophagy and autophagy

Comparative proteomic analysis revealed a panel of downregulated proteins in HepG2D140A mutant cells relative to HepG2WT cells, indicating a severe disruption of cellular waste clearance (autophagy) and compromised mitochondrial homeostasis (Figure 3A). Specifically, key proteins critical for macroautophagy and chaperone-mediated autophagy, including ATG4B, ATG5, ATG12, TBK1, LAMP2 and FKBP8, were significantly depleted. In addition, a significant decrease was detected in a group of lysosomal proteins in cells lacking PRDX6 iPLA2 activity (Figure 3B). Taken together, these findings suggest that these cells have reduced capacity to clear waste and manage the transport of materials destined for autophagy.

**Figure 3.**
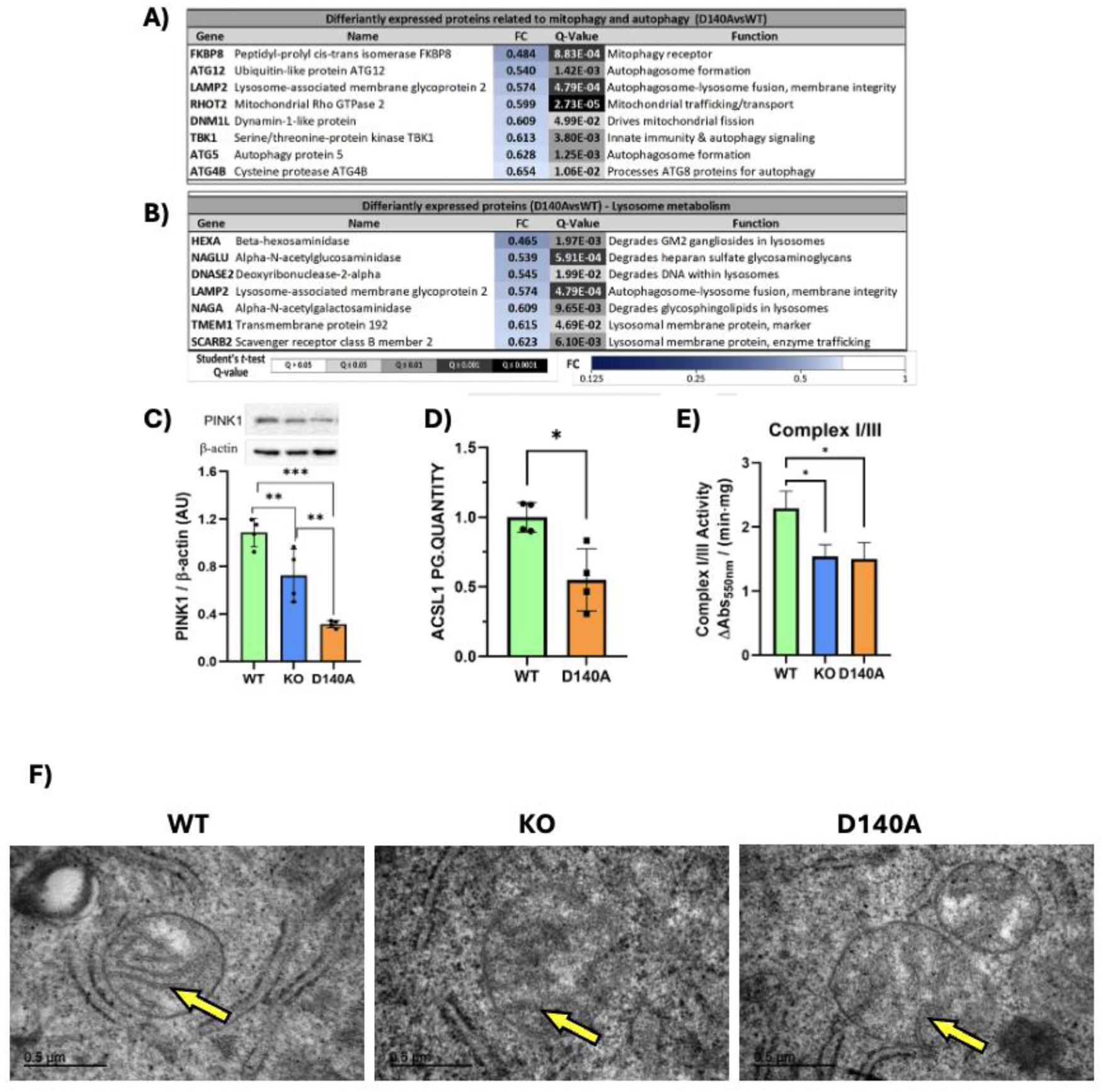
Depletion of PRDX6 iPLA2 activity impairs autophagy, mitophagy, and mitochondrial function and structure in HepG2 cells. (A) Differentially expressed proteins related to mitophagy and autophagy, and (B) lysosomal metabolism (D140A vs. WT) identified by proteomic analysis. (C) PINK1 protein expression quantified by Western-blot in HepG2WT, KO and D140A. (D) ACSL1 protein levels quantified by proteomic analysis in HepG2WT and D140A cells. (E) Mitochondrial Complex I/III activity in HepG2WT, KO and D140A cells. (F) Transmission electron microscopy (TEM) images showing HepG2WT, KO and D140A mitochondrion with cristae (yellow arrows). Student’s *t*-test was used; significance levels: \**P* < 0.05; \*\**P* < 0.01; \*\*\**P* < 0.001.

On the other hand, the downregulation of DNM1L and RHOT2 indicates impaired mitochondrial dynamics by halting fission and intracellular trafficking. No elevation in ROS levels was observed, presumably because D140A mutant cells sustained their peroxidase activity and concurrently exhibited upregulation of alternative antioxidant enzymes, as previously outlined.

These results are fully consistent with the down-regulation of the mitophagy marker PINK1 (Figure 3C), as demonstrated by Western blot. Furthermore, these data corroborate the decreased abundance of ACSL1 (Figure 3D), protein observed in our proteomic analysis, which is known to regulate PINK1-mediated mitophagy. In addition, complex I/III activities were markedly decreased (Figure 3E), presumably due to their vulnerability to oxidative stress and the possible damage to mitochondrial membranes by the presence of certain peroxidized lipids. In this sense, transmission electron microscopy micrographs revealed that mitochondria exhibited an internal disorganization of the mitochondrial cristae compared to WT (Figure 3F). Taken together, these findings indicated that the HepG2D140A cells maintain a delicate balance by activating compensatory mechanisms to sustain survival and prevent cell death.

### Lipidomic analysis demonstrates that iPLA2 deficiency triggers a collapse of lipid homeostasis

The aim of the lipidomic study was to compare the effects of the different genotypes (WT, KO and D140A) on lipid metabolism to identify potential targets or functions associated with the phospholipase A2 activity of PRDX6. A Principal Component Analysis (PCA) model only including the cell samples was generated to identify patterns among the different experimental groups (Figure 4A). As can be observed, the three cell experimental groups clustered separately, according to both the first (t [1]) and the second (t [2]) components of the PCA model (explaining 36.9% and 32.8% of the variability between samples, respectively.

**Figure 4.**
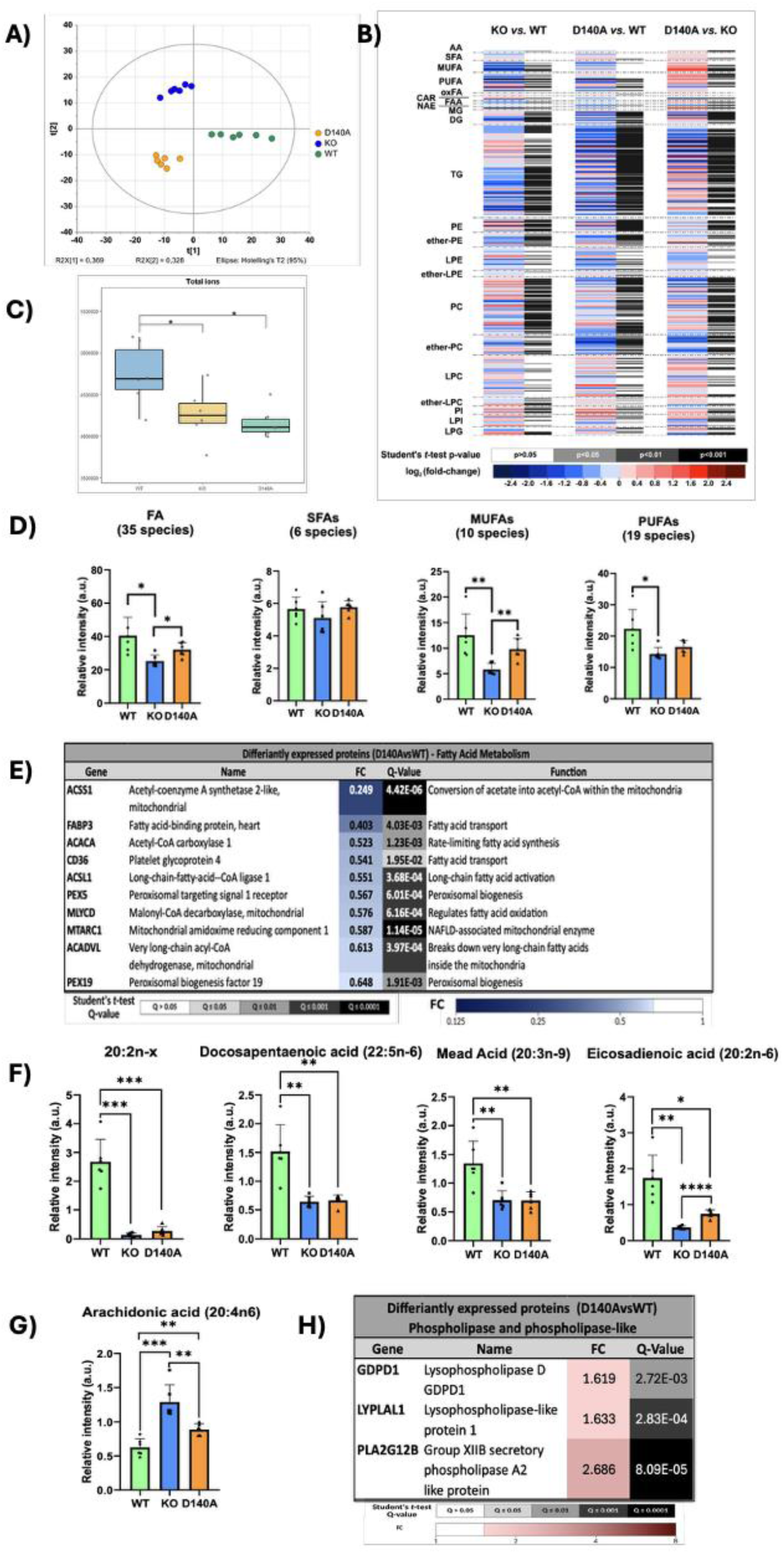
Lack of PRDX6 iPLA2 activity alters the lipidomic profile, disrupts PUFA homeostasis, and triggers a compensatory phospholipase response. (A) Score scatter plot of the Principal Component Analysis (PCA) of HepG2WT, KO and D140A samples. The ellipse represents 95% confidence interval according to Hotelling’s T^2^ test. (B) Heatmaps representing binary unpaired comparisons between KO vs. WT, D140A vs. WT and D140A vs. KO samples, from left to right. Heatmap color codes for log2 (fold-change) and Student’s t-test p-values are indicated at the bottom of the figure. (C) Boxplot showing the Total Ion Current (TIC), representing the sum of all individual sample signals as an indicator of total lipid content. (D) Relative intensity (a.u.) of total FAs, SFAs, MUFAs and PUFAs in HepG2WT, KO and D140A cells. (E) Differentially expressed proteins related to fatty acid metabolism (D140A vs. WT) identified by proteomic analysis. (F) Relative intensity (a.u.) of PUFAs that are decreased upon PRDX6 iPLA2 loss. (G) Relative intensity (a.u.) of arachidonic acid levels in HepG2WT, KO and D140A cells. (H) Differentially expressed proteins related to phospholipase activities (D140A vs. WT) identified by proteomic analysis. Student *t*-test was performed. Significance levels: \**P* < 0.05; \*\**P* < 0.01; \*\*\**P* < 0.001; \*\*\*\**P* < 0.0001.

The obtained results are summarized as a heatmap showing the outcomes of each individual comparison (Figure 4B). As can be observed, plenty of significant differences were found between the three experimental groups assessed. This could be expected due to the notable variances apparent in the multivariate data analysis described above. Of note, the comparisons KO vs. WT and D140A vs. KO showed mostly opposite trends, suggesting that D140A and WT samples are more alike, while those knock-out for PRDX6 are the ones presenting more differences. Nevertheless, D140A vs. WT comparison still highlighted a myriad of significant differences, which were clearly dependent on iPLA2 activity.

Although the analysis performed was semi-quantitative, the Total Ion Current (TIC) of each sample was determined individually. This can give an idea of the total amount of lipids per 2x10^6^ cells. It was observed that the TIC decreased in the same proportion in KO and mutant cells compared to WT cells (Figure 4C).

Described below are the significant changes in specific classes of lipids induced by the loss of phospholipase A2 activity of PRDX6 in hepatocarcinoma HepG2 cells, correlating with the changes identified in our proteomic analysis and align with previously described results.

#### 1. Fatty acids

In this lipidomic study, several saturated fatty acids (SFA) as well as both mono- and poly-unsaturated fatty acids (MUFA and PUFA, respectively) have been assessed in the target cell samples. All significant differences observed are summarized in Figure 4D.

Most of the changes observed in HepG2KO cells consisted of decreases in SFA levels, which could be linked to the lack of iPLA2 activity. Although the D140A mutated cells exhibited partial increases by comparison, these levels did not consistently reach those of HepG2WT cells. This was particularly evident regarding PUFA, for which D140A cells phenocopied KO cells, indicating that these changes might be solely due to the loss of iPLA2 activity, which selectively releases these species. This reduction in PUFA may also be attributed to the oxidation of these compounds, which renders them undetectable using current lipidomic methods, or to an alteration in the *de novo* synthesis of these fatty acids.

The widespread decrease in SFA in HepG2D140A cells aligned with the results obtained from the proteomic analysis (Figure 4E). We found the concurrent downregulation of key proteins that disrupt exogenous fatty acid influx and intracellular trafficking (CD36, FABP3), de novo lipogenesis (ACACA), and mitochondrial acetate activation (ACSS1). Furthermore, lipid catabolism is heavily impaired due to the loss of peroxisomal biogenesis factors (PEX5, PEX19), which compromises very-long-chain fatty acid (VLCFA) oxidation (72). This is accompanied by defects in long-chain fatty acid activation (ACSL1) (73) and mitochondrial β-oxidation (ACADVL) (74). Additionally, the decline in MLYCD potentially elevates malonyl-CoA levels, further obstructing mitochondrial fatty acid entry via CPT1 inhibition. These data demonstrated that mutant cells undergo a collapse of lipid homeostasis, characterized by the concomitant impairment of both anabolic and catabolic lipid pathways.

Despite the significant decrease in free PUFAs in D140A cells (Figure 4F), there was a surprising increase in arachidonic acid (figure 4G). This may be due to increased levels of alternative phospholipases such as LYPLAL1, GDPD1 and PLA2G12B, as observed in the proteomic analysis of the D140A mutant (Figure 4H). On the other hand, it has been described that inhibition of calcium-independent phospholipase A2 prevents arachidonic acid incorporation and phospholipid remodeling in P388D1 macrophages ^66^.

#### 2. Glycerolipids

With respect to TG and DG, both classes of glycerolipids were identified and analyzed, revealing varying trends in the comparisons (Figures 5A and 5B, respectively). Most of the changes consisted of decreases in both the KO and D140A cells compared to the WT controls. In particular, a significant decrease is observed in PUFA-containing TGs and DGs. These results were consistent with the findings for SFA but clearly contrasted with previous findings in another HCC cell line, SNU475, in which TG, particularly those containing long-chain unsaturated fatty acids, were elevated in PRDX6-deficient cells ^5^.

**Figure 5.**
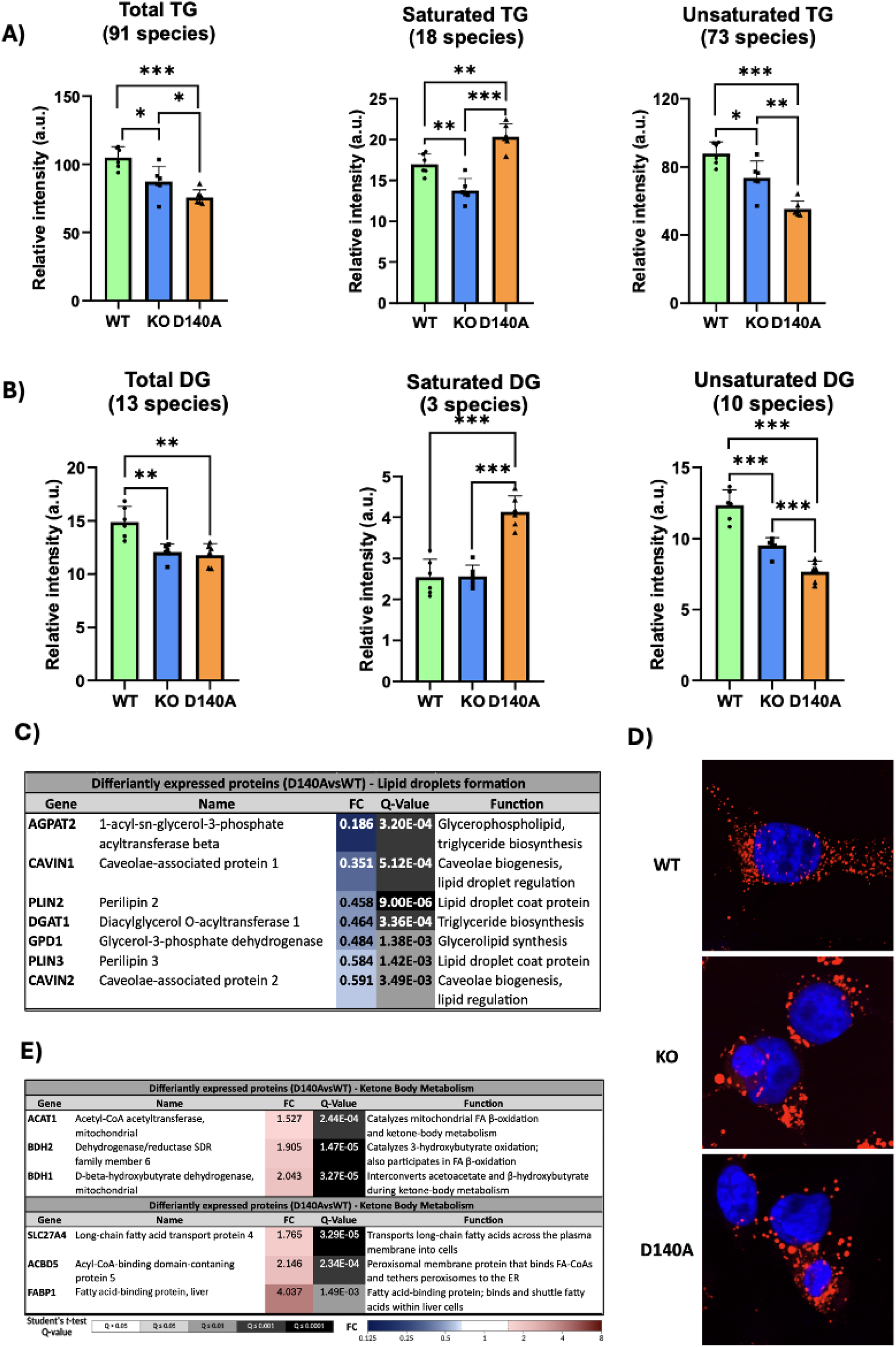
Lack of PRDX6 iPLA2 activity alters glycerolipid levels and lipid droplet biogenesis in HepG2 cells. **(A)** Relative intensity (a.u.) of total TG, saturated TG and unsaturated TG in the three experimental conditions assessed. **(B)** Relative intensity (a.u.) of total DG, saturated DG and unsaturated DG in the three experimental conditions assessed. **C)** Differentially expressed proteins related to lipid droplets biogenesis (D140A vs. WT) identified by proteomic analysis. **(D)** Nile red staining of HepG2WT, KO and D140A cells. Lipid droplets are stained red, and the nuclei are stained blue. **(E)** Differentially expressed proteins related to lipid metabolism (D140A vs. WT) identified by proteomic analysis. Student *t*-test was performed. Significance levels: \**P* < 0.05; \*\**P* < 0.01; \*\*\**P* < 0.001.

Diglycerides and triglycerides are the core lipid building blocks required for LD biogenesis ^67,68^. This metabolic process occurs at the ER membrane to safely store metabolic energy and protect the cell from the lipotoxic effect of free fatty acid. Proteomic analysis of HepG2D140 cells revealed significant disruptions in LD biogenesis, storage, and metabolism (Figure 5C). The downregulation of key enzymes, including GPD1, DGAT1, and AGPAT2, suppresses triglyceride synthesis, while downregulation of CAVIN1, CAVIN2, PLIN2, and PLIN3 impairs cell signaling, lipid regulation, and LD stabilization. Visually, fluorescence microscopy images of the three HepG2 cell types stained with Nile Red confirmed these alterations, showing a reduced number of lipid droplets and increased size heterogeneity in both knockout and mutant cells compared to the WT (Figure 5D). Nevertheless, we observed a significant upregulation of a specific enzymes, including ACAT1, BDH2, BDH1, SLC27A4 (FATP4), ACBD5 and FABP1 (Figure 5E), indicating an enhancement in lipid trafficking, fatty acid β-oxidation, and ketone body metabolism which represents a desperate metabolic shift. These results point to a disruption of normal LD storage and structure while simultaneously driving a compensatory shift toward membrane remodeling, increased fat oxidation, and ketone body production to manage cellular energy demands as a consequence of the loss of PRDX6 iPLA2 activity.

#### 3. Glycerophospholipids

The comparisons analyzed revealed numerous significant differences in glycerophospholipid levels among WT, KO, and D140A cells. Due to the substrate specificity of PRDX6 iPLA2 activity toward distinct phospholipid species, the trends observed in phospholipids containing AA at the *sn*-2 position were analyzed independently as potential targets. The levels of AA were drastically decreased in all the glycerophospholipid classes (Figure S1), but with the striking exception of three phosphatidylcholine (PC), two phosphatidylethanolamine (PE) and two phosphatidylinositol (PI) containing species which may be indicative of specificity of the phospholipase A2 activity of PRDX6 for these lipids with AA at sn2 position (Figure 6A).

**Figure 6.**
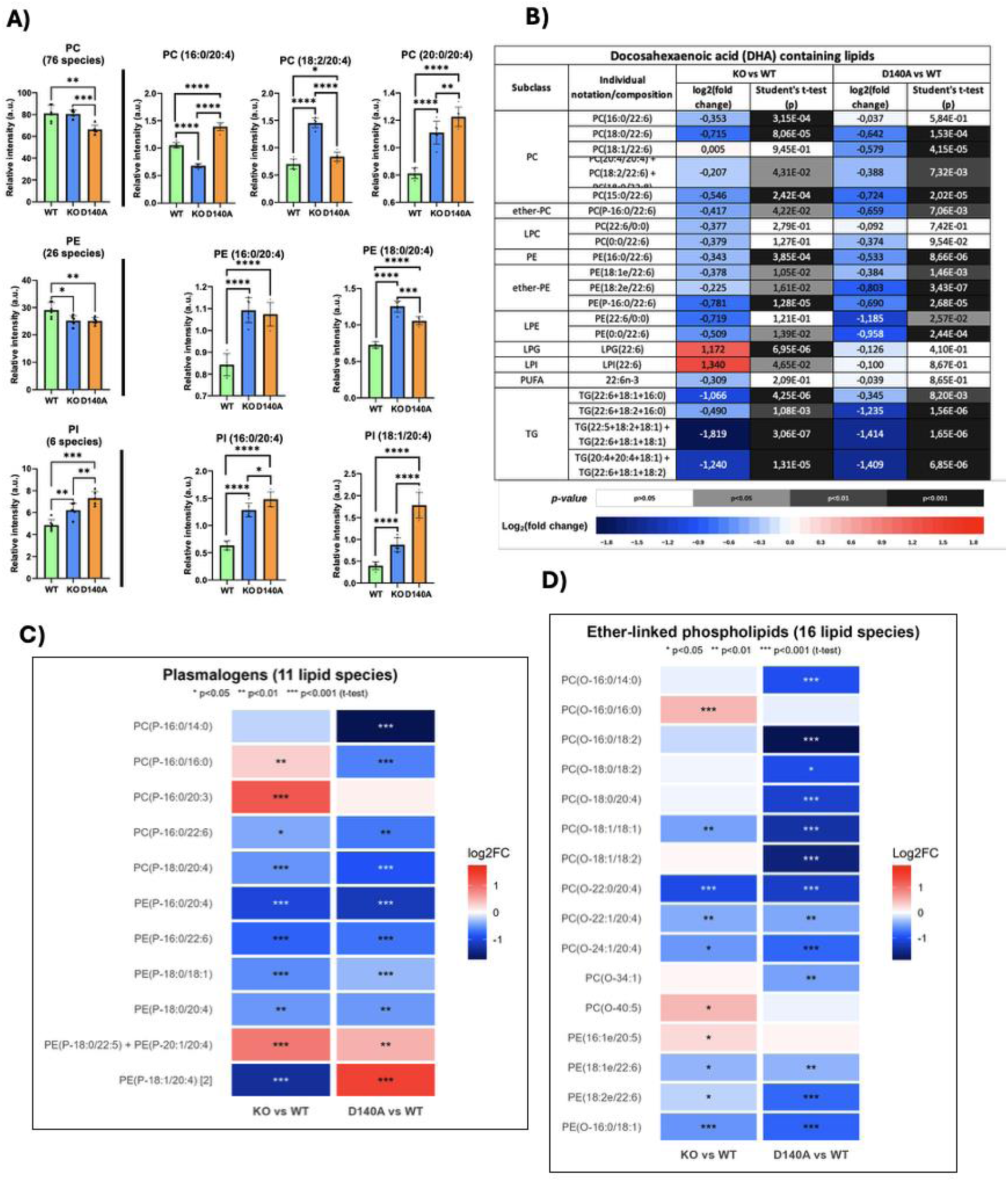
PRDX6 iPLA2 activity deficiency remodels arachidonic acid- and docosahexaenoic acid-containing phospholipids and downregulates ether-linked glycerophospholipids in HepG2 cells. (A) Relative intensity (a.u.) of total PC, PE and PI, and of selected arachidonic acid (20:4)-containing species of each class, in HepG2WT, KO and D140A cells. (B) Docosahexaenoic acid (DHA, 22:6)-containing lipids in KO and D140A cells relative to WT. Blue/red sections denote lipids that were reduced or increased in WT vs KO and D140A vs KO, respectively; gray/black bars indicate significant changes. (C) Heatmaps showing the comparisons of KO vs. WT (left) and D140A vs. WT (right) for plasmalogens and (D) ether-linked glycerophospholipids. Colors represent log2 fold-change, and asterisks indicate Student’s *t*-test *p*-values. Significance levels: \**P* < 0.05; \*\**P* < 0.01; \*\*\**P* < 0.001.

In addition, our results demonstrated that all docosahexaenoic acid (DHA) containing glycerophospholipids decreased drastically in both KO and D140A cell groups compared to WT, except free DHA, which did not change (Figure 6B). These decreases were consistent with a previous lipidomic analysis conducted on SNU475 cell line lacking PRDX6 ^5^. Despite significant differences found in the lipid profiles of HepG2 and SNU475 cells, characterized by their differing degrees of dedifferentiation, in the case of DHA, the profile resulting from the absence of PRDX6 is identical to and consistent with that observed in HepG2D140A cells; therefore, iPLA2 activity must be responsible for this generalized decrease in DHA.

Another group of glycerophospholipids that was severely affected by the loss of PRDX6 iPLA2 activity comprises ether-linked glycerophospholipids. A significant majority of these species (32 in total), including both plasmanyl (ether-linked) and plasmenyl (vinyl-ether-linked) forms, were markedly reduced in both HepG2KO and D140A cells (Figure 6C and 6D). Notably, within the plasmenyl subclass, only two species exhibited a clear increase in D140A cells: PE(P-20:1/20:4) and PE(P-18:1/20:4). Both represent canonical plasmalogens possessing AA esterified at the sn-2 position. Similar results were obtained in the SNU475 cell line, which lacked PRDX6 ^5^. These data demonstrate that loss of PRDX6 or its iPLA2 activity in HepG2 cells severely disrupts plasmalogen homeostasis, which likely underlies the signaling, metabolic, and cell-cycle alterations previously documented in PRDX6-devoid cells. Furthermore, the strong correlation between PRDX6 deficiency and the accumulation of AA-containing plasmalogens in D140A mutant cells suggests that these specific lipid species may serve as substrates for the iPLA2 enzymatic activity of PRDX6. Plasmalogens (especially ethanolamine plasmalogens) are highly enriched in AA at the sn-2 position, and the specific cleavage of this position by plasmalogen-selective iPLA2 represents a major pathway for AA release.

### PRDX6 iPLA2 activity deficiency disrupts function and inter-organelle communication among peroxisomes, ER, mitochondria, and lipid droplets

The peroxisomes, ER, mitochondria, and LDs cooperate extensively to regulate lipid metabolism and drive or protect against ferroptosis.

Peroxisomes synthesize ether lipids, specifically plasmalogens, that play critical roles as endogenous antioxidants and mediators of membrane dynamics and they have been related to ferroptosis ^69^. Our lipidomics results showed a significant decrease in ether lipids resulting from the lack of phospholipase activity. These findings are fully supported by the proteomics data, which revealed the downregulation of several proteins essential for proper peroxisomal function (Figure 7A). For instance, we observed a drop in peroxisomal AGPS expression related to the reduction of ether lipid production. Furthermore, we observed a decrease in proteins related to peroxisomal biogenesis and matrix protein import (PEX5 and PEX19), as well as those associated with peroxisomal metabolism (AGXT, AGPS, PIPOX and MLYCD) and mitochondria-peroxisome communication (DNM1L and RHOT2). Therefore, the coordinated decrease in these specific proteins indicates a severe impairment of peroxisomal biogenesis, metabolic flux, organelle dynamics, and signaling pathways, potentially contributing to cellular stress and ferroptosis.

**Figure 7.**
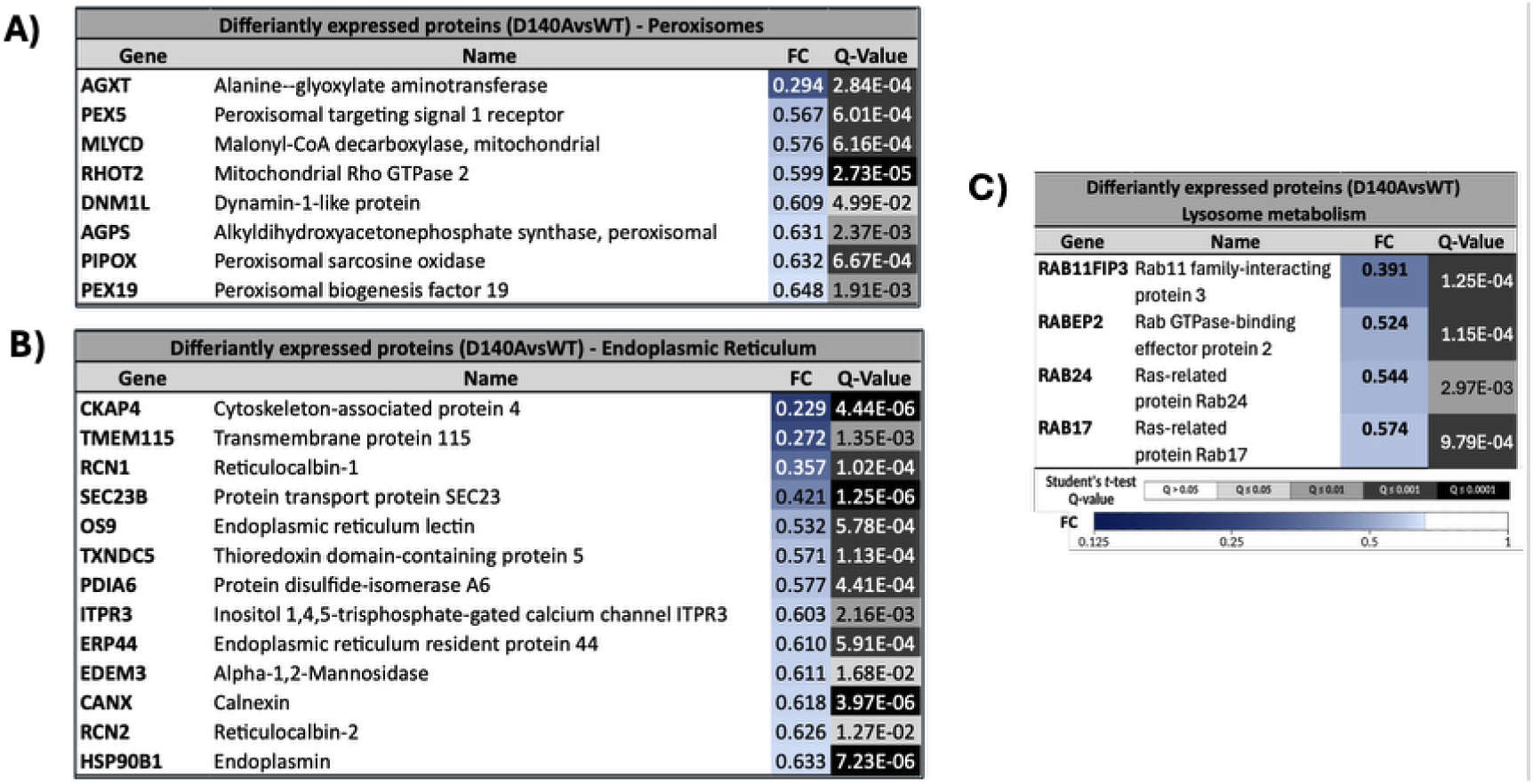
Loss of PRDX6 iPLA2 activity disrupts peroxisome and endoplasmic reticulum protein homeostasis. (A) Differentially expressed peroxisome related proteins in HepG2 D140A vs HepG2 WT cells; (B) Differentially expressed endoplasmic reticulum (ER) proteins in HepG2 D140A vs HepG2 WT cells. Corrected Student’s t-test q-values obtained by proteomic analysis are shown.

On the other hand, our findings in HepG2D140A cells demonstrated a significant downregulation of ACSL1 (Figure 4C) and DGAT-1 (Figure 5C), two key enzymes involved in triglyceride synthesis. This suppression may trigger a cascade of cellular alterations that ultimately drive the cells toward ferroptotic cell death. The ACSL1 activates long-chain fatty acids to facilitate triglyceride synthesis and subsequent storage within lipid droplets, while concurrently regulating PINK-mediated mitophagy ^70^. These results are consistent with those shown in Figure 3, which revealed a significant decrease in mitophagy markers in HepG2D140A cells compared to HepG2WT cells. Similarly, DGAT-1, an ER enzyme, catalyzes the synthesis of triglycerides from diacylglycerol and acyl-CoA for lipid droplet storage. Depletion of DGAT-1 blocks mitophagy, induces lipotoxicity, and ultimately triggers ferroptotic cell death ^71^. Furthermore, the downregulation of specific ER-associated proteins, namely HSP90B1, RCN2, CANX, EDEM3, ERP44, ITPR3, PDIA6, TXNDC5, OS9, SEC23B, RCN1, TMEM115, and CKAP4 (Figure 7B) indicated a severe collapse of the ER proteostasis network ^72^. Finally, in our study, we observed alterations in LP in D140A and KO cells when compared to WT as we have described in Figure 5. The expression of Rab and Rab-related proteins such as RAB24, RABEP2, RAB17 and RAB11FIP3, which are involved in LD homeostasis, was decreased in D140A mutant (Figure 7C). These proteins interact with LD to regulate the formation of membrane contact sites between intracellular membranes of organelles, such as the mitochondria, and to control lipid trafficking and homeostasis ^73^.

Taking all these results into account, it becomes evident that the lack of PRDX6 iPLA2 activity induces a breakdown of lipid homeostasis characterized by the simultaneous impairment of organelles such as peroxisomes, ER, mitochondria and LDs, making cells susceptible to ferroptotic death.

## DISCUSSION

In human hepatocelular carcinoma, malignant progression, metabolic reprogramming, and evasion of cell death are critically dependent on redox maintenance and lipid homeostasis ^74^. To investigate PRDX6’s iPLA2 role in these processes, this study utilized wild-type (WT), PRDX6-knockout (KO), and iPLA2-inactive mutant (D140A) HepG2 cell lines. The D140A mutation drastically lowered iPLA2 activity to match KO levels, a phenotype successfully replicated in WT cells using the inhibitor MJ33, confirming its utility for *in vivo* applications.

PRDX6’s iPLA2 activity hydrolyzes the sn-2 bond of glycerophospholipids, generating free fatty acids and lysophospholipids, and its deficiency severely disrupts the cellular lipidome by preventing proper phospholipid recycling. This disruption alters membrane fluidity, curvature, and stability, ultimately reducing cell viability. Additionally, the lack of liberated fatty acids depletes lipid-derived secondary messengers (e.g., eicosanoids), altering enzyme expression. While prior research mostly focused on iPLA2β and iPLA2γ ^75–77^; this study provides the first comprehensive proteomic, lipidomic, and ferroptosis cell death analysis of PRDX6 iPLA2 deficiency in hepatocellular carcinoma.

Our findings demonstrate that, while certain phenotypes depend strictly on the complete absence of PRDX6, others are due exclusively to the selective loss of either iPLA2 activity or peroxidase activity. Dissecting this, PRDX6 lack increases intracellular ROS, but the iPLA2 D140A mutant maintains wild-type ROS levels because its peroxidase activity remains intact. Conversely, lipid peroxides significantly increase in D140A cells compared to WT, though less severely than in KO cells. This intermediate elevation occurs because the intact peroxidase activity reduces dangerous phospholipid hydroperoxides to stable alcohols ^78^ but without iPLA2 activity, these structurally altered lipids cannot be cleaved or recycled, remaining trapped and nonfunctional within the membrane ^79^. Furthermore, there must also be oxidized lipids that cannot be released due to the lack of PRDX6 iPLA2 activity, and which the peroxidase activity is unable to reduce.

In addition, our results demonstrate that both enzymatic activities of PRDX6 are also necessary to sustain cell proliferation. When both activities are absent, hydrogen peroxide (H_2_O_2_) and lipid peroxides impair cell cycle progression, consistent with previous studies showing G2/M phase arrest and decreased migratory signaling in PRDX6-deficient HCC cell lines ^56,57^. The iPLA2 activity of PRDX6 is known to trigger downstream mitogenic signaling, such as the activation of the extracellular signal regulated kinase (ERK) and p38 mitogen-activated protein kinase (MAPK) pathways, by releasing arachidonic acid for eicosanoid synthesis ^80^. It has also been reported that the two activities of PRDX6 work together to promote the development of lung tumors through the CCL5-associated, PRDX6-mediated activation of the JAK2/STAT3 pathway ^81^. This gradient in proliferative impairment demonstrates that neither enzymatic activity is completely redundant; rather, they exert distinct yet complementary control over cell cycle progression.

A pivotal finding of this study is that both PRDX6 KO and D140A mutant cells exhibit a similar, heightened sensitivity to ferroptosis induced by erastin, and significant accumulation of intracellular ferrous iron (Fe^2+^). Ferroptosis is an iron-dependent form of regulated cell death driven by the accumulation of lipid hydroperoxides within plasma and organellar membranes. The finding that D140A mutant cells fully phenocopy the KO cells regarding ferroptosis sensitivity demonstrates that the iPLA2 activity of PRDX6 is strictly required for ferroptosis resistance, independently of its peroxidase domain integrity. Although the peroxidase activity of PRDX6 can directly reduce phospholipid hydroperoxides, its iPLA2 activity is specifically responsible for hydrolyzing oxidized fatty acyl chains at the *sn-2* position of glycerophospholipids, allowing the subsequent re-acylation with native fatty acids to completely regenerate membrane integrity. In the absence of functional iPLA2 activity, oxidized fatty acids or those reduced by peroxidase and whose structure is no longer functional cannot be excised from the lipid bilayer. This structural failure leads to an accumulation of oxidized lipids, which we confirmed is more elevated in D140A mutant cells than in WT cells. This compromised state is further exacerbated by a uniform downregulation of glutathione peroxidase 4 (GPX4) in D140 mutant and KO cells. Because GPX4 is the primary enzymatic defender against ferroptosis, its downregulation creates a severe vulnerability to erastin. To counteract this deficit, HepG2D140A cell line orchestrates a compensatory upregulation of the standard glutathione synthesis and recycling machinery, alongside other mitochondrial antioxidant proteins. However, this antioxidant response is clearly insufficient to overcome the loss of membrane lipid editing normally afforded by PRDX6. Additionally, the loss of iPLA2 function matches the KO phenotype in altering iron handling proteins compared to WT cells, with the resulting iron dysregulation driving the increased ferroptotic flux.

PRDX6 is fundamental for mitochondrial function, as its deficiency severely impairs mitochondrial quality control and mitophagic flux. Specifically, both HepG2KO and D140A cells show a downregulation of PINK1 and LC3B, correlated with decreased levels of the regulatory proteins ACSL1 and ATG4B, which cripples the canonical mitophagic pathway for degrading damaged mitochondria ^82^. This results in dysfunctional mitochondrial accumulation with reduced complex I/III activities and structurally abnormal cristae, observed by electron microscopy, due to lipid peroxidation. Furthermore, the D140A mutant exhibits a downregulation of proteins critical for macroautophagy, chaperone-mediated autophagy, and lysosomal function, halting autophagosome biogenesis and leading to cytotoxic debris accumulation ^83–86^.

Despite mitochondrial dysfunction, ROS levels did not increase. This is attributed to the cells sustaining their peroxidase activity and upregulating alternative antioxidant enzymes.

Therefore, although HepG2D140A cells manage to maintain a fragile homeostatic balance to survive, these defects leave them highly vulnerable to external stressors, specifically making them prone to erastin-induced ferroptosis.

Lipidomic and proteomic analyses confirm that loss of PRDX6 iPLA2 activity causes a collapse in lipid homeostasis, characterized by decreased total lipid content, reduced free fatty acids (especially PUFAs), and altered glycerophospholipid profiles. Both KO and D140A cells show a severe depletion of arachidonoc acid (AA) and docosahexaenoic acid (DHA) containing species, likely due to systematic lipid peroxidation or synthesis defects. While most PUFA modifications (elongation and desaturation) take place in the endoplasmic reticulum, the final steps required to complete the synthesis of DHA (and alternatively the retroconversion of certain AA metabolites) specifically involve peroxisomal β-oxidation. We have demonstrated the impairment of peroxisomal function, evidenced by the downregulation of key proteins involved in ether lipid synthesis and peroxisomal biogenesis. Additionally, the endoplasmic reticulum, which is the primary site for the synthesis of phospholipids and the incorporation of PUFAs into membrane lipids, also shows signs of dysfunction. PUFAs synthesized by the ER and incorporated into phospholipids and neutral lipids are either transferred to the mitochondria to regulate energy production or channeled into lipid droplets to mitigate lipotoxicity and oxidative damage (90). The endoplasmic reticulum (ER), mitochondria, and lipid droplets (LDs) form a strictly regulated tripartite network. Our data demonstrate that the regulation of this tripartite network is disrupted; in addition to alterations in the ER, we have also observed mitochondrial damage, both functional and structural, and disorganization in the biogenesis of lipid droplets.

The significant accumulation of AA-containing lipids at the sn-2 position in KO and D140A cells suggests these are specific substrates for the PRDX6 iPLA2 activity, which can no longer cleave and release them. Unexpectedly, free arachidonic acid levels also increase in these cell lines. This anomaly indicates a disruption in the continuous deacylation/reacylation cycle (the Lands cycle) that normally keeps intracellular free AA low. The resulting excess of non-esterified AA acts as a toxic mediator, ultimately inducing cell death ^87^.

Therefore, iPLA2 activity of PRDX6 is a master regulator of lipid homeostasis in HCC. Lacking it changes the physical structure of membranes and deprives the cell of critical lipid signaling molecules. Many transcription factors act as “lipid sensors.” For example, PPARs (Peroxisome Proliferator-Activated Receptors) are activated by free fatty acids, and SREBPs (Sterol Regulatory Element-Binding Proteins) respond to membrane lipid composition. When iPLA2 fails to produce these lipid ligands, these transcription factors cannot bind to DNA, shutting down or over-activating the expression of downstream metabolic enzymes ^88^. The cell senses this crisis and drastically reprograms its gene expression to compensate, affecting other enzymatic pathways. The cell is actively trying to bypass the broken classical pathways by shifting its reliance toward alternative lipid utilization, membrane remodeling, and ketone body production, as evidenced by upregulation of enzymes involved in these pathways, to sustain survival. However, the systemic disruption of essential trafficking and organelle biogenesis factors creates an insurmountable metabolic bottleneck, resulting in a decoupled phenotype, with significant consequences for the survival of cancer cells and their susceptibility to ferroptosis.

## CONCLUSION

Here, we report the first comprehensive proteomic, lipidomic, and ferroptotic pathway analysis detailing the molecular effects driven by PRDX6 iPLA2 activity deficiency in hepatocellular carcinoma (HCC). This study provides definitive evidence that peroxidase and iPLA2 activities of PRDX6 exert both overlapping and highly distinct regulatory controls in HCC. We demonstrate that PRDX6 iPLA2 activity is crucial for lipid homeostasis, cell proliferation, resistance to ferroptosis, and proper organelle function in HepG2 cells. Its loss leads to profound metabolic and structural changes, partially compensated by upregulation of alternative antioxidant and metabolic pathways, but ultimately increases cellular vulnerability to stress and death by ferroptosis. Inhibiting this specific domain successfully mimics the total knockout by triggering iron accumulation, suppressing mitophagy, and opening the floodgates to ferroptotic cell death, while bypassing full systemic toxicities. In conclusion, targeting the iPLA2activity of PRDX6, which can be achived using MJ33, may represent a highly viable therapeutic strategy for hepatocelular carcinoma.

## Supporting information

Lipidomic dataset

## DATA AND CODE AVAILABILITY

All data supporting the findings of this study are available within the article or its supplementary materials. The proteomics data have been deposited in the ProteomeXchange Consortium via the PRIDE partner repository with the dataset identifiers: PROJECT ID: PXD086432. Token: jWy41izbXYEl.

## AUTHOR CONTRIBUTION

C. Alicia Padilla: Conceptualization, Supervision, Funding acquisition, Validation, Project administration, Writing —review and editing. Raquel Requejo-Aguilar: Conceptualization, Formal analysis, Supervision, Funding acquisition, Project administration, Writing — review and editing. Ángel Ortiz-Alcántara: Software, Formal analysis, Validation, Investigation, Methodology. Alberto Ortiz-Olivencia: Software, Formal analysis, Investigation, Visualization, Methodology. Raúl González-Ojeda: Conceptualization, Supervision, Validation, Investigation, Methodology. Brian McDonagh: Conceptualization, Formal analysis, Supervision, Writing.

## ACKNOWLEDGEMENTS

Technical support by the staff of the Microscopy and Proteomics facilities, SCAI, University of Cordoba and IMIBIC, and the company Rubió Metabolomics (Bizkaia, Spain) are acknowledged.

## STAR+METHODS

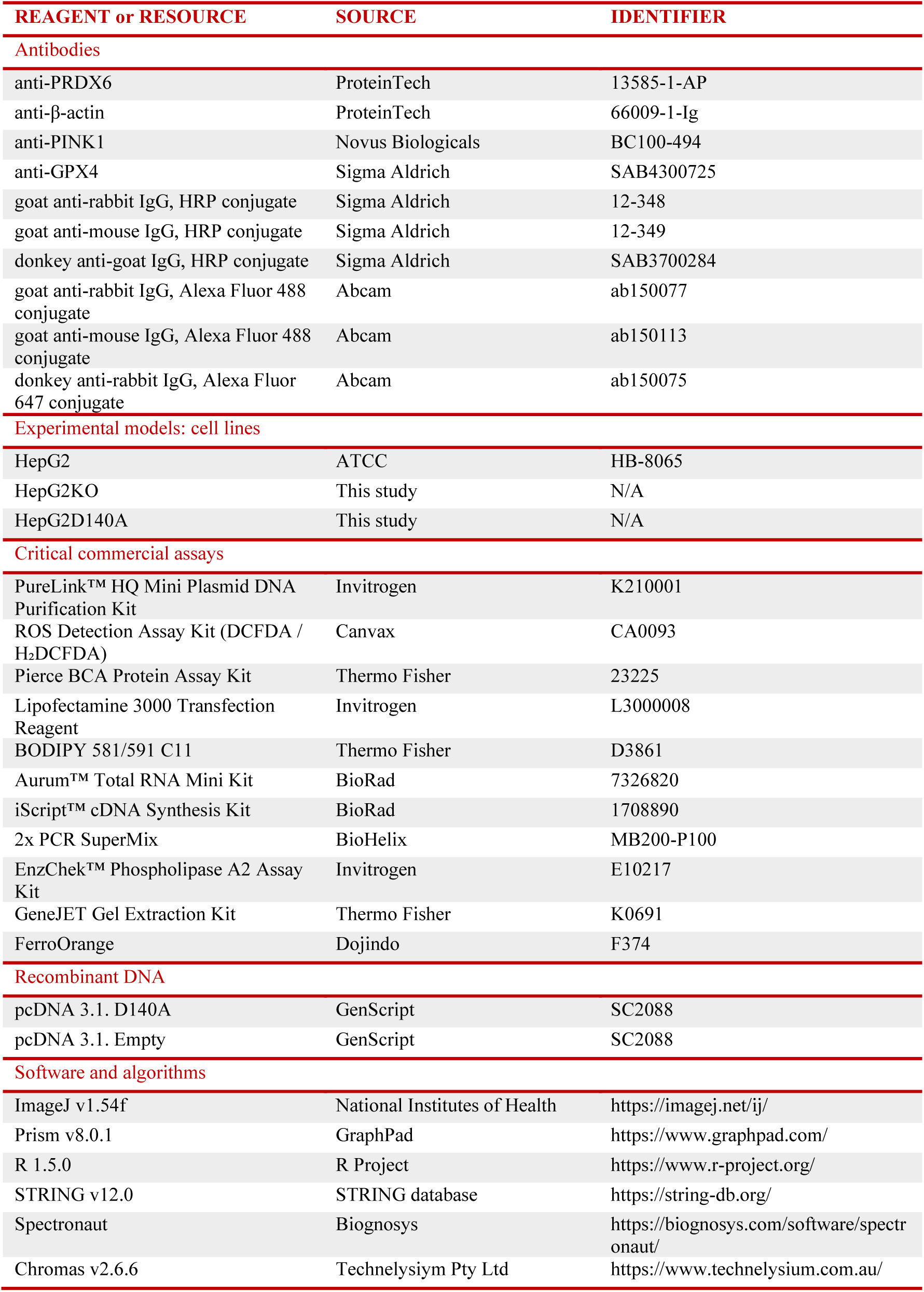
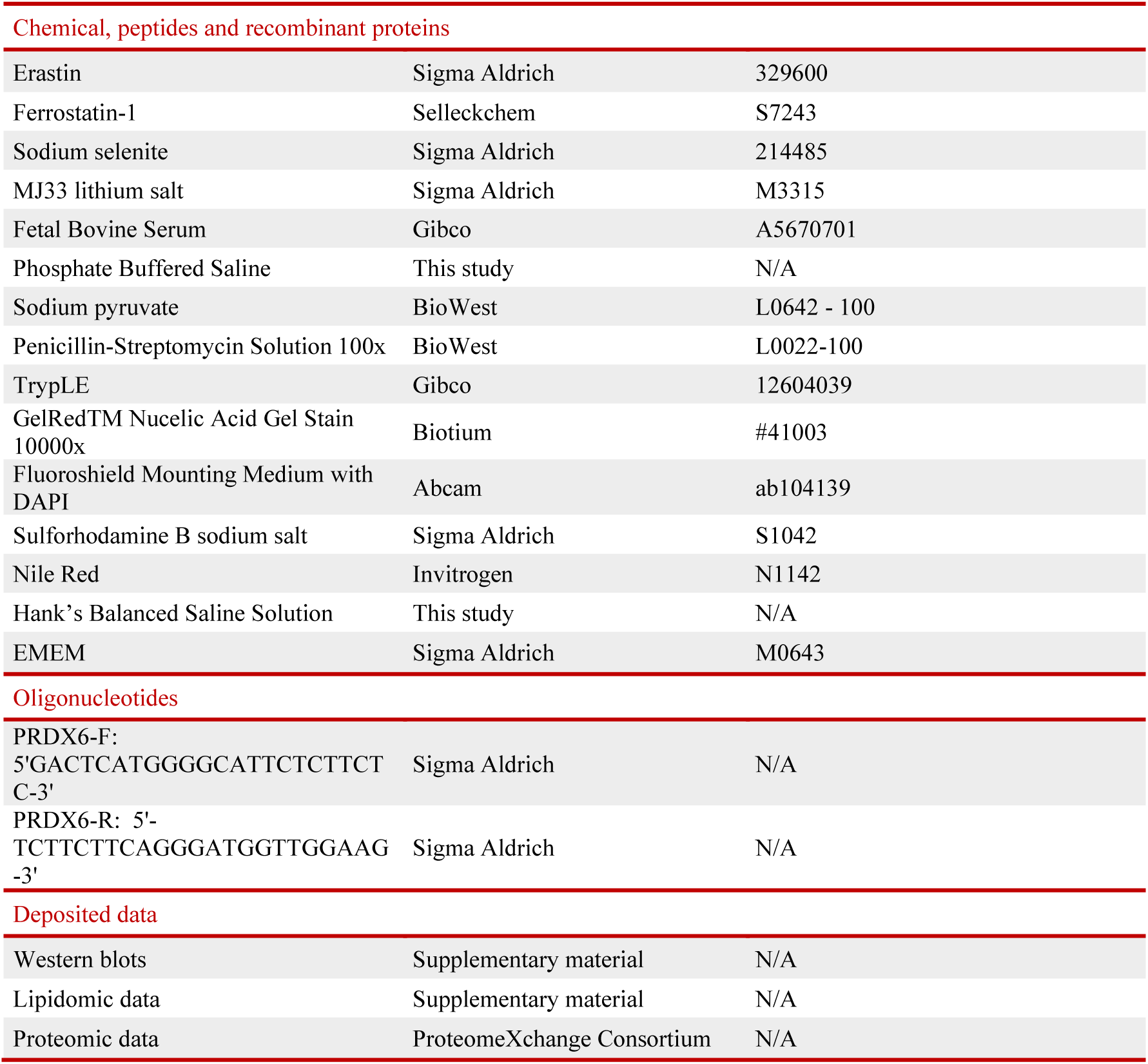

## METHODS

### Cell lines and culture conditions

HepG2 standard cell line was obtained from American Type Culture Collection (ATCC) and it was used as a positive control (HepG2WT or WT). The HepG2 lacking PRDX6 (HepG2KO or KO) cell line was generated by using CRISPR/Cas9 technology as previously described ^56^ and used both, as a negative control and the backbone for the construction of the HepG2 cell line lacking iPLA2 activity (HepG2D140A or D140A). All cell lines were grown in aseptic conditions and pH 7.4 in an incubator with a 5% CO_2_ atmosphere and 37°C using Eagle’s Minimal Essential Medium (EMEM) supplemented with 10% fetal bovine serum (FBS, Gibco), 1mM sodium pyruvate (Biowest) and 100 U/L penicillin, 100 μg/mL streptomycin, 0.25 μg/mL amphotericin. For obtention of cell lysates, cells were either detached by trypsinization (TrypLE, Biowest) or scrapped with lysis buffer, according to the requirements.

### HepG2D140A cell line generation

HepG2D140A cell line was generated by transfecting HepG2KO cell line with the pcDNA3.1. plasmid containing PRDX6D140A cDNA, and posterior enrichment, selection and validation. 5 × 10^5^ cells were seeded in a 6-well plate and were transfected for 6 h upon reaching 80% of confluence. Transfection was performed using the lipofectamine 3000 kit according to manufacturer guidelines, using 2.5 µg of plasmid DNA and a ratio of 1:2:2 [DNA (µg): P3000 reagent (µL): lipofectamine (µL]. Transfected cells were then treated sequentially with increasing concentrations of H_2_O_2_ (50, 100, 150, 200 and 250μM) every time medium was refreshed as an enrichment treatment for two weeks. A limit dilution of the surviving cells was then performed by seeding 1.5 cells/well in 96-well plates for the isolation of clones and let to grow. Clones were analyzed for PRDX6 by Western blot using specific antibodies. Presence of mutant HepG2D140A in analyzed clones was validated by Sanger DNA sequencing.

### cDNA synthesis and sequencing

Total RNA was isolated from 2 × 10⁶ cells using the Aurum™ Total RNA Mini Kit (Bio-Rad), following the manufacturer’s instructions. Up to 1,000ng of total RNA was used for cDNA synthesis with the iScript™ cDNA Synthesis Kit (Bio-Rad). PCR amplification was performed using 1000ng of cDNA as template, the PRDX6-F and PRDX6-R primers, and PCR SuperMix(BioHelix). The amplification was carried out for 30 cycles consisting of denaturation at 95°C for 15 s, annealing at 55°C for 15 s, and extension at 72°C for 30 s. A total of 10 μL of the PCR product was loaded on a 1.2% (w/v) agarose gel containing 0.01% (v/v) GelRed® (Biotium) and electrophoresis was performed at 80V for 45 minutes. DNA bands were visualized under UV light, excised from the gel, and purified using the GeneJET Gel Extraction Kit (Thermo Scientific). Purified PCR products were sequenced on an ABI Prism 3130XL Genetic Analyzer (Applied Biosystems) at the Genomics Service of the Central Research Support Service (SCAI), University of Córdoba and analyzed using Chromas Software (v2.6.6).

### Assays for the enzymatic activity calcium-independent phospholipase A2

PLA2 activity in HepG2 cells was measured in 96-well plates using the EnzChek Phospholipase A2 Assay (Invitrogen) with Red/Green BODIPY at pH 7.0 without calcium. Cells were lysed via sonication in PLA2 buffer (50 mM Tris-HCl, 1 mM EGTA, pH 7.0, containing protease and phosphatase inhibitors) and centrifuged at 15,000 × g for 15 min. The resulting supernatant was harvested, and 50 μg of protein was preincubated at 37 °C for 10 min, both with and without 10 μM MJ33 (10 μM) ^89^. After adding 50 μL of the Red/Green BODIPY substrate, fluorescence was monitored for 1 h using a Varioskan microplate reader (Thermo Scientific) at excitation/emission wavelengths of 470/515 nm.

### Mitochondrial Complex I and III activity assay

Cells were pelleted and lysed in 20 mM phosphate buffer (pH 7.5) containing 1 mM MgCl₂, 1 mM EDTA, and 1 mM PMSF. Mechanical lysis was performed using micropestles for 1 min, and lysates were subsequently incubated for 10 min at 4°C. Samples were centrifuged at 800 × g for 5 min at 4°C, and supernatant was collected. NADH-cytochrome *c* oxidoreductase activity was measured spectrophotometrically at 550nm in 50mM phosphate buffer (pH 7.5) containing 100μM cytochrome c, 80 μM KCN, and 100 μM NADH, in the presence or absence of 100 μM rotenone. Absorbance at 550 nm was recorded every 10 s for 3 min, both before and after the addition of NADH, at 30°C. Enzymatic activity was normalized to protein concentration. Complex I activity was calculated as the rotenone-sensitive fraction, while Complex III activity corresponded to the rotenone-insensitive fraction.

### Cell proliferation determination

HepG2WT, HepG2KO and HepG2D140A cells (1.5 x 10^4^) were seeded in 96 well plates, treated as required after 24 hours, and allowed to grow for 1-4 days. Cells were then collected and counted, generating the corresponding growth curves. Additionally, cell proliferation was measured using SRB assay, as described in Vichai, V *et al.* (2006) ^90^.

### Cell lysis, protein extracts preparation, SDS-PAGE and Western-blot

Cells were lysated at 4°C with RIPA lysis buffer (NaH_2_PO_4_ 15 mM, Na_2_EDTA 10 mM, NaCl 158 mM, SDS 0.1% v/v, Triton X-100 1% v/v) containing protease inhibitors (SIGMA) and NaF 1mM as phosphatase inhibitor (SIGMA), and supernatants were collected. Protein concentration was determined using a BCA Assay Kit (Thermo Fisher Scientific) with BSA as the standard. SDS–PAGE was performed in 10-15% acrylamide gels and electrophoresis was carried out in a Mini-protean Tetra Cell system (Bio-Rad) for 90 min at 120 V, using “NZYColour Protein Marker II, 10-245kDa” as molecular ladder, and loading 15 ug of protein. After electrophoresis, proteins were transferred to a nitrocellulose membrane using a wet transfer system for 80 min at 100V. Transfer and protein load were checked by staining with Ponceau reagent. Membranes were blocked with 5% non-fat milk in Tris-buffered saline, 0.1% v/v Tween-20 detergent (TBS-T) for 1 h at room temperature and then incubated overnight at 4 °C with gentle agitation with the corresponding dilutions of primary antibodies (**Table 1**). Membranes were then washed three consecutive times with TBS-T and incubated with the corresponding peroxidase-conjugated secondary antibody (**Table 1**) for 1 h at room temperature. For detection of chemiluminescent signal, membranes were added “NZY Advanced ECL” (NZYTech, Lisbon, Portugal) solution and images were taken in a ChemiDoc image analyzer (Bio-Rad Laboratories; Hercules, CA, USA) and quantified by densitometry with ImageJ software version 1.54f (National Institutes of Health, Bethesda, MD, USA), using actin, GAPDH or Ponceau staining as a reference for loading normalization.

### ROS assay and quantification

Determination of ROS levels in HepG2WT, HepG2KO and HepG2D140A cells was carried out using the commercial DCFDA/H_2_DCFDA ROS Assay Kit (Canvax, Córdoba, Spain). 1.5x10^4^ cells were seeded in 96-well plates and allowed to grow for 48 hours. Then, cells were incubated at 37°C with H_2_DCFDA 50µM for 1 hour protected from light, washed with commercial assay buffer and treated with 100µM H_2_O_2_ for 1 hour. Fluorescence was determined in Varioskan LUX Fluorimeter λex = 485 nm; λex = 530 nm.

### Lipid Peroxidation

Lipid peroxidation levels were quantified using the fluorescent probe BODIPY C11 581/591 (Invitrogen). 2x10^5^ cells were seeded in 12-well plates and allowed to grow for 24h, treated with erastin (4 µM) and ferrostatin-1 (10 µM) and incubated 48 h. Cells were then incubated with BODIPY C11 581/591 2.5 µM in HBSS for 30 minutes and washed twice with HBSS. Cells were harvested and centrifuged for 5min at 300 *g*. Supernatant was discarded and cell pellet was resuspended in 500 µL PBS. Fluorescence was measured through flow cytometry using BD Accuri C6 Plus flow cytometer (BD Biosciences).

### Intracellular iron measurement

1.5x10^4^ HepG2WT, HepG2KO and HepG2D140A cells were seeded in 96-well plate and allowed to grow for 48 hours. Then, cells were washed twice with PBS and incubated with FerroOrange (Dojindo) 1 µM in PBS for 30 minutes at 37°C and 5% CO_2_. Fluorescence was measured using Varioskan LUX Fluorimeter (ThermoFisher Scientific) at λex = 561 nm; λex = 600 nm.

### Inmunofluorescence

HepG2WT, HepG2KO and HepG2D140Acells (8x10^4^) were seeded in coverslips coated with Poly-D-Lys and allowed to grow overnight with complete cell culture medium. Cells were fixed with 4% paraformaldehyde (PFA) in PBS for 20 minutes at room temperature. Permeabilization was conducted with 0,3% Triton X-100 in PBS for 10 minutes at room temperature and washed 10 minutes twice with PBS. Lipid droplets staining was performed adding 1 µM Nile Red solution in PBS and incubating for 90 minutes at room temperature. PRDX6 detection was performed using anti-PRDX6 antibody (Proteintech) 1:1000 in 2% BSA w/v in PBS overnight at 4°C and anti-IgG rabbit Alexa Fluor 647 nm in 2% BSA w/v in PBS for 90 min. Cells were then washed 10 minutes twice with PBS and preparations were mounted using mounting medium with DAPI. Preparations were imaged by employing a conventional fluorescence microscope Leica DM6B (Wetlzar, Germany) and analyzed with FIJI software.

### Global proteome analysis

Cells were lysed at 4°C in RIPA lysis buffer (15 mM NaH₂PO₄, 10 mM Na₂EDTA, 158 mM NaCl, 0.1% v/v SDS, 1% v/v Triton X-100) supplemented with protease inhibitors (Sigma) and 1 mM NaF as a phosphatase inhibitor (Sigma), and supernatants were collected. Protein concentration was determined using a BCA Assay Kit (Thermo Fisher) with BSA as standard, and samples were normalized to 3 μg/μL. Four biological replicates per experimental group were analyzed. Protein digestion was performed by addition of 10 μL of 12.5 ng/μL trypsin solution (Promega, WI, USA) in 25mM NH₄HCO₃ and incubated at 37°C overnight, before being stopped by addition of 3 μL of trifluoroacetic acid (1.5% final concentration). Digested samples were dialyzed through Pierce detergent removal columns to eliminate any remaining traces of detergent and dried in a SpeedVac. Proteolytic digestion, sample preparation, and LC-MS/MS analyses were performed at the Proteomics Facility (SCAI), University of Córdoba. Peptides were separated on a nano-LC system using an IonOpticksAurora analytical column (25cm) with a 30-minute gradient and analyzed in DIA-PASEF mode on a Bruker timsTOF mass spectrometer, using an ion mobility (1/K0) window of 0.6–1.6 Vs/cm². Raw data were processed using Spectronaut software against the UniProtKB/Swiss-Prot curated human protein database (June 2026 release), using a library-free (directDIA) search strategy. A protein was considered significantly expressed if identified in at least three of four replicates in at least one experimental group. Differential expression of proteins was assessed using log2 fold-change and Q-value cutoffs of 0.6 and 0.05, respectively. Functional enrichment analyses were performed using STRING, KEGG/GO pathway annotations, and the Reactome pathway database. Additionally, predicted upstream regulators were identified using the DoRothEA consensus regulon network (R package/decoupleR), considering regulons with confidence levels A–C.

### Global Lipidome Analysis

The lipidomic profiling was conducted at Rubió Metabolomics (Bizkaia, Spain) in accordance with specific company guidelines. Cell samples from 6 different experiments (2x10^6^) were harvested, frozen in liquid nitrogen and stored at -80°C until shipment. Metabolite extraction was performed by fractionating cell samples into a pool of species sharing similar physicochemical properties, employing methanol/chloroform as a solvent system. First, cell pellets were resuspended with water. Protein precipitation was induced by adding methanol enriched with internal standards, followed by vortexing. Subsequently, methanol/chloroform enriched with internal standards and pure chloroform were added to the mix. Samples were incubated at - 20°C for 30 minutes and collected for each platform.

Two distinct UHPLC-MS platforms were employed for the optimal profiling of:

- Fatty acids, bile acids, steroids and lysoglycerophospholipids: Supernatants were collected, dried under vacuum and reconstituted in methanol prior to transfer to plates for UHPLC-MS analysis.
- Glycerolipids, glycerophospholipids, sterol lipids and sphingolipids: Water was added to 500 uL of extract, and samples were incubated again for 30 minutes at -20°C. Samples were centrifuged at 18 000 × *g* for 15 min at 4°C to facilitate phase separation. The organic layer was collected, dried under vacuum, resuspended in acetonitrile/isopropanol (1:1) and transferred to vials for UHPLC-MS analysis.

Additionally, different types of quality control (QC) samples were used to assess data quality. For each analytical platform, randomized sample injections were performed, with QC calibration and validation extracts uniformly interspersed throughout the entire batch run. Chromatographic separations were achieved using ACQUITY UPLC systems (Waters Corp., Milford, USA) across all platforms. An LCT Premier XE Time-of-Flight (ToF) (Waters Corp.) and a Xevo G2 QTof (Waters Corp.) mass spectrometer was used for Lipidomics Platforms 1 and 2, respectively. All data were processed using the TargetLynx application manager for MassLynx 4.1 software.

Normalization factors were calculated for each metabolite by dividing its intensity in each sample by the recorded intensity of an appropriate internal standard in that same sample, following the procedure described by Martínez-Arranz et al. ^91^. Multivariate data analysis was performed via principal components analysis (PCA), where the data matrix was reduced to a series of principal components (PCs), each representing a linear combination of metabolite peak areas. Additional statistical evaluations performed were the Shapiro–Wilk test, Student’s t-test, Wilcoxon test, and Volcano plot (see Appendix 1 for further details).

### Transmission electron microscopy

Preparation of samples for transmission electron microscopy was carried out at the SCAI of the University of Córdoba. 2 ×10^6^ cells were harvested and fixed with 2.5% v/v solution of glutaraldehyde in 0.1 mol/l phosphate buffer, pH 7.4. Cells were fixed for 30 min in a 1% osmium tetroxide solution in the same buffer, dehydrated using increasing concentrations of ethanol, and embedded in Epon resin. 90 nm sections were cut using an ultramicrotome, stained with 5% uranyl acetate and 5% lead citrate, and analyzed on a JEM1400 transmission electron microscope at 80 kV.

### Statistical analysis

Results are expressed as mean ± SD of at least three independent experiments (N=3), each one with at least three technical replicas (n=3-10). Statistical analysis of the data was performed by unpaired Student’s *t*-test or one/two-way ANOVA, depending on the experiment design, as detailed in the figure legends. Levels of significance considered are as follows: \**p*-value<0,05; \*\**p*-value<0,01; \*\*\**p*-value<0,001; \*\*\*\**p*-value<0,0001.

