## Supplementary figures and images for "The Phospholipase A2 Activity of Peroxiredoxin 6 Protects Hepatocarcinoma Cells Against Ferroptosis and Mitophagy Impairment"

### Figure S1

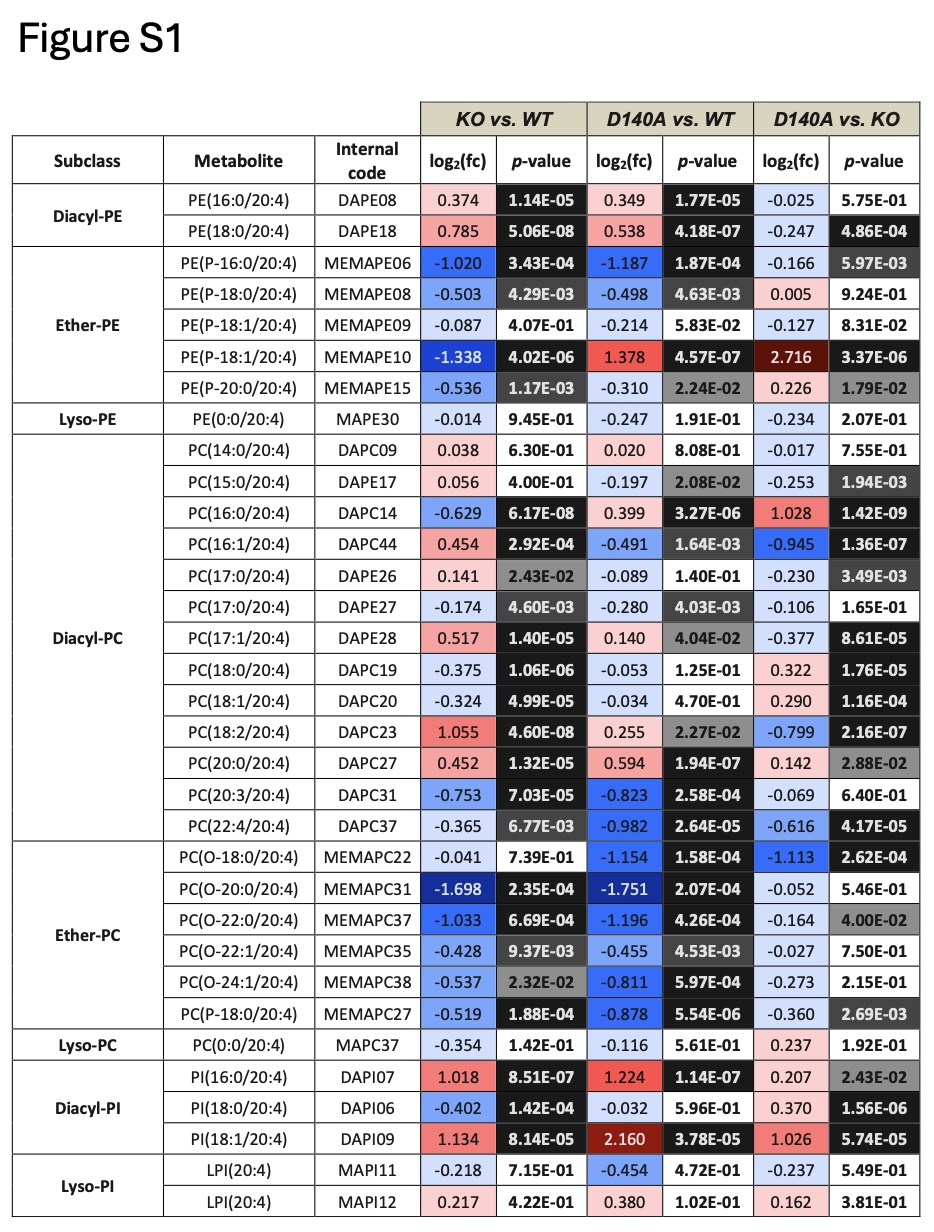
